# The first dinosaur egg remains a mystery

**DOI:** 10.1101/2020.12.10.406678

**Authors:** Lucas J. Legendre, David Rubilar-Rogers, Alexander O. Vargas, Julia A. Clarke

## Abstract

A recent study by Norell et al. (2020) described new egg specimens for two dinosaur species, identified as the first soft-shelled dinosaur eggs. The authors used phylogenetic comparative methods to reconstruct eggshell type in a sample of reptiles, and identified the eggs of dinosaurs and archosaurs as ancestrally soft-shelled, with three independent acquisitions of a hard eggshell among dinosaurs. This result contradicts previous hypotheses of hard-shelled eggs as ancestral to archosaurs and dinosaurs. Here we estimate the ancestral condition for dinosaur and archosaur eggs by reanalyzing the original data from Norell et al. and that from a recent study on reptile eggshells (Legendre et al., 2020) with the addition of these new dinosaur specimens. We show that the recovery of dinosaur eggs as ancestrally soft-shelled is conditioned by the discretization of a continuous character (eggshell thickness), the exclusion of turtle outgroups from the original sample, and a lack of branch length information. When using a larger sample, calibrated trees, and a definition of hard-shelled eggs referencing their unique prismatic structure, we recover dinosaur and archosaur eggs as either hard-shelled or uncertain (i.e. equal probability for hard- and soft-shelled). This remaining ambiguity is due to uncertainty in the assessment of eggshell type in two dinosaur species, i.e. ∼1% of the total sample. We conclude that more reptile egg specimens and a strict comparative framework are necessary to decipher the evolution of dinosaur eggs in a phylogenetic context.

---

In a recent study, Norell et al.^1^ describe remarkable egg specimens for two species of nonavian dinosaurs. Using spectroscopic and statistical analyses, the authors identify these new specimens as soft-shelled, their shell being described as a completely unmineralized, proteinaceous egg membrane. This represents an exceptional discovery in the study of dinosaur reproduction^2^: all previously known dinosaur eggs, including those of extant birds, are hard-shelled, i.e. they present a mineralized calcareous layer that is made of column-shaped calcium carbonate structures called shell units^3^, on top of the egg membrane. Hard-shelled eggs are by far the most common egg type among extant reptiles – also found in all crocodilians, turtles, and some gekkos^2^ – while soft-shelled eggs, lacking shell units, are found in other lepidosaurs^4,5^. Norell et al.^1^ perform an ancestral state reconstruction of eggshell structure on a phylogeny of 112 extant and extinct reptiles, using both maximum likelihood (ML) and maximum parsimony (MP), and find soft-shelled eggs to be the ancestral condition in both dinosaurs and archosaurs. This result challenges the previously accepted view of hard-shelled eggs being ancestral for these two groups^2^, and implies that dinosaurs independently acquired hard-shelled eggs at least three separate times. We agree with the conclusions of the authors that these new specimens represent the first described dinosaurian soft-shelled eggs, and the hypothesis that the first dinosaur egg was soft has important implications for the reproductive strategies of this emblematic group of reptiles. However, in our own analyses^6^, published on the same day, we found support for the traditional hypothesis. Was this due to only to the impact of the two new eggs, different taxonomic sampling more generally, or how soft-shelled eggs were defined? Here, we reassess the ancestral condition for dinosaur and archosaur eggs, with reanalysis of both the original data used to propose this hypothesis^1^ and our own dataset^6^ with the addition of these new dinosaur specimens^1^ (Figs 1, 2).

**Figure 1.**
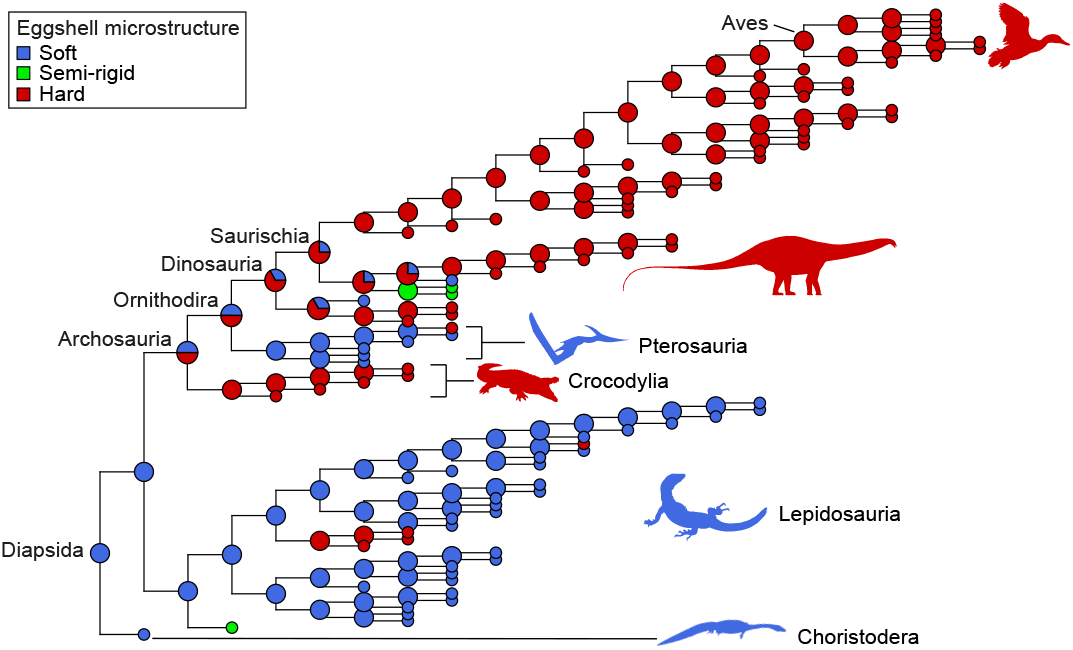
Ancestral state reconstructions for eggshell structure using the tree and matrix of Norell et al.^1^ recover dinosaurs as ancestrally hard-shelled if the scoring of only two previously published taxa is modified (*Lufengosaurus* and *Massospondylus* from soft to semi-rigid; see text). Maximum Parsimony ASR (MP; using asr_max_parsimony in ‘castor’^11^) is shown; for ML, see Extended Data Figure 1.

**Figure 2.**
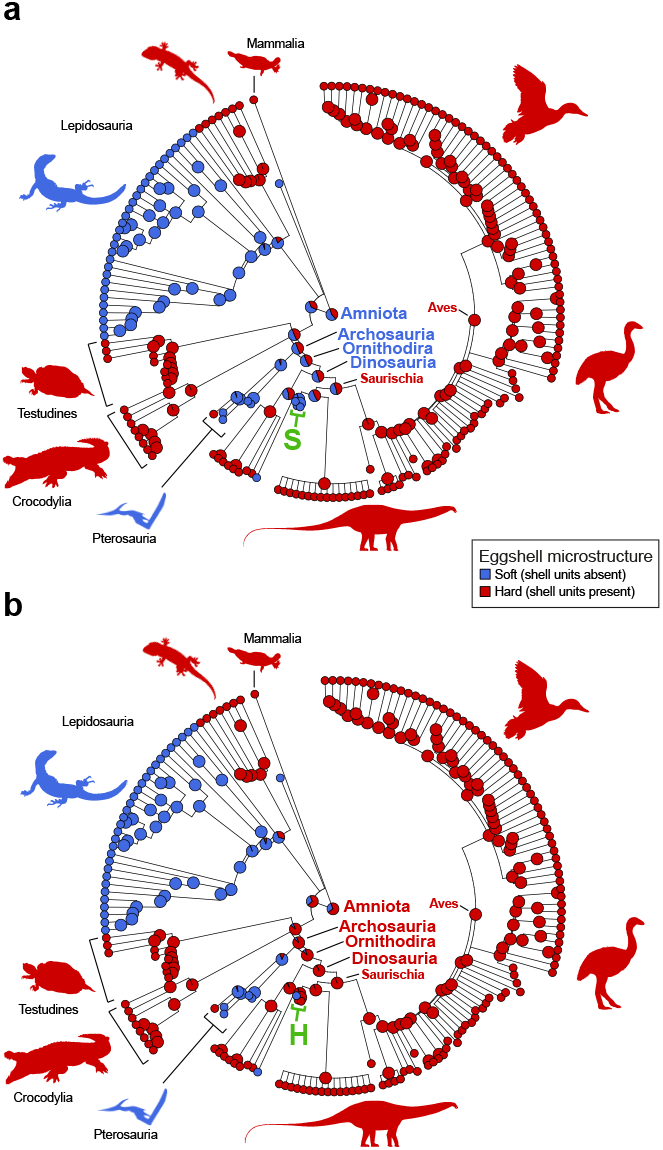
Ancestral state reconstructions for eggshell structure after adding the new fossils to Legendre et al.^6^. Dinosaurs as either ambiguously soft- or hard-shelled with the scorings of Norell et al. (a) or hard-shelled with our scoring (b). Again, the difference in optimization is due not to the new eggs, but depends on the coding for the two dinosaurs *Lufengosaurus* and *Massospondylus* – coded as soft-shelled (green “S”) in a^1^, and as hard-shelled due to the presence of mineralized shell units^6^ (green “H”) in b; analyses used make.simmap in ‘phytools’^15^.

Key to note, these two studies^1,6^ do not use the same definition of “soft-shelled”. Reptile eggshells have been classified in many ways, some contradictory. Some reptile eggs (e.g. turtles) present a folded eggshell that superficially looks soft, but has mineralized shell units similar in structure and ontogeny to those of hard-shelled eggs (e.g. birds and crocodiles)^3–5^. Such eggshells are sometimes considered intermediate between hard and soft, and have been referred to as “pliable” or “semi-rigid”^1,5,7^. The approach used by Norell et al. to distinguish hard-shelled and soft-shelled eggs^1^ did not reference the presence or absence of shell units, but used three categories derived from the ratio “calcareous layer thickness/total eggshell thickness”. The range of values for this ratio was subdivided into three states (soft/semi-rigid/hard shell), with respective thresholds of 0.5 and 0.67 between them^1^. They cited two references^7,8^ in support of this practice. Neither of those references, however, actually defines such thresholds. Both studies^7,8^ discuss reptilian eggshells in a taxonomic context, describing three general eggshell types (soft/semi-rigid/hard) and hypothesizing they might represent different ontogenetic strategies and gas exchange solutions. However, these authors^7,8^ acknowledge further work is required to assess if those qualitative, “somewhat artificial” categories correspond to meaningful differences (Packard et al.:142)^8^. These thresholds may be able to be derived empirically, e.g. using gap weighting methods, but there is currently no consensus on the relative accuracy of such procedures in a phylogenetic context^9^.

Identifying a soft-shelled egg category by thresholds of thickness, versus presence/absence of mineralized shell units, impacts the reconstructed ancestral states for dinosaurs. Two sauropodomorph dinosaurs, *Lufengosaurus* and *Massospondylus*, were scored as soft-shelled^1^, but their eggshells present mineralized shell units^10^ (unlike the newly-reported eggs). A change in discretized character state from soft to semi-rigid or hard for only these two taxa out of 112 (replicating the original MP analysis^1^ in R using ‘castor’^11^) results in dinosaurs recovered as ancestrally hard-shelled, and archosaurs recovered as ambiguous (either soft- or hard-shelled; Fig. 1). Recovery of dinosaur eggs as ancestrally soft-shelled is thus highly sensitive to small single assumptions for small sets of taxa. The exclusion in the original analysis^1^ of the proposed sister-group to archosaurs, turtles, is also expected to affect these outcomes.

Recovery of the ancestral dinosaur egg as soft-shelled is also dependent on branch length assumptions. The original ancestral state reconstruction (ASR) analyses^1^ did not include any time-calibrated branch length information (Extended Data Fig. 1a), which is a well-known significant source of bias for ancestral state reconstruction^12,13^. Mesquite was used^1^, and when branch length information is not provided, Mesquite ML ASR automatically assigns a default length of 1.0 to all branches^14^, resulting in a tree with unrealistic branch lengths; MP ASR does not consider branch length information^12^. An identical topology with different default branch lengths (e.g. basic ultrametricized tree; Extended Data Fig. 1b), as well as MCMC stochastic character mapping in ‘phytools’^15^ (ER model, 1000 simulations), considered in recent literature to be more accurate than ML in this context^12,13^, return archosaurs and dinosaurs as ancestrally hard-shelled with reptiles as ancestrally soft-shelled, in agreement with previous scenarios for eggshell evolution^2^.

In our recent study of a giant Antarctic soft-shelled egg^6^, we distinguish hard and soft eggshells on presence or absence of mineralized shell units (see above), and not based on their relative thickness. Our ASR of reptile eggshell thickness, using a time-calibrated tree and including turtles, recovered shell units in gecko eggs as non-homologous to those of other reptiles, as previously suggested^4^, but hard-shelled eggs as ancestral to dinosaurs and archosaurs. However, we did not include the new eggs nor several soft-shelled pterosaur eggs sampled in Norell et al.^1^ Adding these new species (total = 178 taxa) renders the recovered ancestral condition for archosaurs and dinosaurs as ambiguously optimized (Fig. 2a). If *Mussaurus* and *Massospondylus* (i.e. ∼1% total sample) are treated as hard-shelled as they would be in our original analysis (based on their preserved evidence of shell units), archosaurs and dinosaur eggs are again recovered as ancestrally hard-shelled (Fig. 2b).

Recovered ancestral eggshell structure for archosaurs and dinosaurs is highly influenced by branch length information, character coding, and taxon sampling. Newly described soft-shelled eggs^1,6^ show that the reproductive strategies of reptiles were much more diverse than previously thought, and highlight that our knowledge of eggshell evolution remains incomplete. The two datasets analysed here do not support dinosaurs as ancestrally laying soft-shelled eggs, as this result requires a discretized continuous character (eggshell thickness), and exclusion of a proposed sister taxon of archosaurs, turtles. If the scoring of only two taxa in the original study is changed, this result is no longer recovered. Other sources of phylogenetic uncertainty, such as imprecise branch length information or low sample size for some clades, limit both recent studies^1,6^. With state-of-the-art techniques and new exceptionally preserved eggshells to be discovered, there is more we have yet to learn about the evolution of reproductive modes in dinosaurs and other reptiles.

## Acknowledgements

The authors thank J. Wiemann, M. Fabbri, and D. Zelenitsky for providing the original character matrix and phylogenetic trees used in their study. Figures include silhouettes edited from PhyloPic (detail in Legendre et al.^6^)

## Data availability

All data used in this paper were taken from published literature cited in the main text. Formatted tables, phylogenetic trees, and R code for all analyses performed or replicated in this paper are available on GitHub at https://github.com/LucasLegendre/norelletal_response.

## Author contributions

LJL and JAC conceived the research; LJL performed the analyses and wrote the paper; all authors discussed and contributed to the final draft of the manuscript.

The authors declare no competing interests.

**Extended Data Figure 1.**
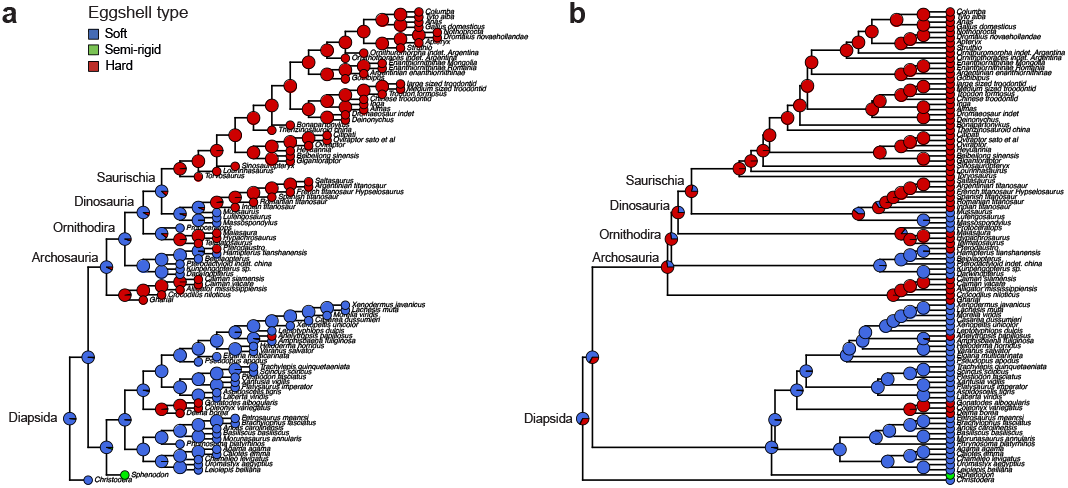
Ancestral state reconstructions (ASR) for diapsid eggshell structure (soft-shelled/hard-shelled), using the tree and matrix of Norell et al.^1^. **a**, ASR using the original tree of Norell et al., with all branch lengths equal to 1. **b**, ASR using an ultrametric version of the tree in **a**, i.e. with an identical topology but different branch lengths. Both ASR were performed using make.simmap in ‘phytools’^15^ (see text).

## References

1. Norell, M. A. et al. The first dinosaur egg was soft. Nature 583, 406–410 (2020).

2. Sander, P. M. Reproduction in early amniotes. Science 337, 806–808 (2012).

3. Mikhailov, K. E. Fossil and recent eggshell in amniotic vertebrates: fine structure, comparative morphology and classification. Spec Pap Palaeontol 56, 1–76 (1997).

4. Choi, S., Han, S., Kim, N.-H. & Lee, Y.-N. A comparative study of eggshells of Gekkota with morphological, chemical compositional and crystallographic approaches and its evolutionary implications. PLOS ONE 13, e0199496 (2018).

5. Schleich, H. H. & Kåstle, W. Reptile egg-shells: SEM atlas. (Fischer, 1988).

6. Legendre, L. J. et al. A giant soft-shelled egg from the Late Cretaceous of Antarctica. Nature 583, 411–414 (2020).

7. Kusuda, S. et al. Diversity in the Matrix Structure of Eggshells in the Testudines (Reptilia). Zool Sci 30, 366–374 (2013).

8. Packard, M. J., Packard, G. C. & Boardman, T. J. Structure of Eggshells and Water Relations of Reptilian Eggs. Herpetologica 38, 136–155 (1982).

9. Bardin, J., Rouget, I., Yacobucci, M. M. & Cecca, F. Increasing the number of discrete character states for continuous characters generates well-resolved trees that do not reflect phylogeny. Integr Zool 9, 531–541 (2014).

10. Stein, K. et al. Structure and evolutionary implications of the earliest (Sinemurian, Early Jurassic) dinosaur eggs and eggshells. Sci Rep 9, 4424 (2019).

11. Louca, S. & Doebeli, M. Efficient comparative phylogenetics on large trees. Bioinformatics 34, 1053–1055 (2018).

12. Joy, J. B., Liang, R. H., McCloskey, R. M., Nguyen, T. & Poon, A. F. Y. Ancestral Reconstruction. PLOS Comput Biol 12, e1004763 (2016).

13. Cunningham, C. W., Omland, K. E. & Oakley, T. H. Reconstructing ancestral character states: a critical reappraisal. Trends Ecol Evol 13, 361–366 (1998).

14. Maddison, W. P. & Maddison, D. R. Mesquite: a modular system for evolutionary analysis. (2019).

15. Revell, L. J. phytools: an R package for phylogenetic comparative biology (and other things). Methods Ecol Evol 3, 217–223 (2012).

